# Smartphone-based ear-EEG to study sound processing in everyday life

**DOI:** 10.1101/2023.01.31.526197

**Authors:** Daniel Hölle, Martin G. Bleichner

## Abstract

In everyday life, people differ in their sound perception and thus sound processing. Some people may be distracted by construction noise while others do not even notice. With smartphone-based mobile ear-electroencephalography we can measure and quantify sound processing in everyday life by analyzing presented sounds and also naturally occurring ones. Twenty-four participants completed four controlled conditions in the lab (1h) and one condition in the office (3h). All conditions used the same paired-click stimuli. In the lab, participants listened to click tones under four different instructions: no task towards the sounds, reading a newspaper article, listening to an audio article, or counting a rare deviant sound. In the office recording, participants followed daily activities while they were sporadically presented with clicks, without any further instruction. In addition to the presented sounds, environmental sounds were recorded as acoustic features (i.e., loudness, power spectral density, sounds onsets). We found task-dependent differences in the auditory event-related potentials (ERPs) to the presented click sounds in all lab conditions, which underline that neural processes related to auditory attention can be differentiated with ear-EEG. In the office condition, we found ERPs comparable to some of the lab conditions. The N1 amplitude to the click sounds beyond the lab was dependent on the background noise, probably due to energetic masking. Contrary to our expectation, we did not find a clear ERP in response to the environmental sounds. Overall, we showed that smartphone-based ear-EEG can be used to study sound processing of well defined-stimuli in everyday life.

## 1 Introduction

How do people perceive and thus process sounds under natural listening conditions in everyday life? It is evident that people differ in their sound perception over time and context. Some sounds are attended, others are not noticed. However, it is entirely unclear how much of their acoustic environment people perceive spontaneously and how much goes unnoticed. To obtain more insights into sound processing, we conducted a smartphone-based mobile ear-electroencephalography (ear-EEG) study in the lab and in everyday life. For this, we used a two-fold approach: we present well-defined stimuli and we record naturally occurring sounds, which we then relate to the ongoing EEG.

First, we are interested in the neural response to well-defined stimuli in an everyday life context. From previous mobile EEG studies (e.g., Debener, Minow, Emkes, Gandras, & de Vos, 2012; Dehais, Roy, & Scannella, 2019; Hölle, Meekes, & Bleichner, 2021; Ladouce, Donaldson, Dudchenko, & Ietswaart, 2019; Reiser, Wascher, & Arnau, 2019; Scanlon, Townsend, Cormier, Kuziek, & Mathewson, 2019; Somon, Giebeler, Darmet, & Dehais, 2022; Zink, Hunyadi, Huffel, & Vos, 2016) we know that auditory attention can be measured beyond the lab. Following this line of work, we used paired click sounds as well-defined stimuli in an otherwise acoustically uncontrolled environment. The paired-click paradigm has been used to study noise annoyance and distractability (Bak et al., 2017; Gjini, Burroughs, & Boutros, 2011; Shepherd, Hautus, Lee, & Mulgrew, 2016; Shepherd, Lodhia, & Hautus, 2019) and allows to study the neural response to a surprising sound (first tone) and to a cued sound (second tone).

As opposed to these aforementioned beyond-the-lab studies in which participants had an explicit task instruction (e.g., count a deviant sound), in this study, we are interested in how people process their everyday life soundscape normally without a specific task instruction on how to behave towards sounds. However, without an overt task instruction, the interpretation of the results is difficult. To grasp how to interpret the beyond-the-lab results, in this study, participants also listened to the same stimuli under controlled lab conditions with specific instructions on how to behave towards the sounds (see methods for detailed instructions). The results of the lab conditions serve as a reference to interpret the data acquired beyond the lab (cf. Ladouce, Mustile, Ietswaart, & Dehais, 2022 for a similar approach).

Second, we are interested in the neural response to naturally occurring sounds. In a previous study, we have shown that it is possible to compute ERPs to onsets extracted from an everyday acoustic scene using a smartphone app (Hölle, Blum, Kissner, Debener, & Bleichner, 2022). Here we take this approach beyond the lab and record all sound onsets that occurred in the participants’ environment. With this data, we can, on the one hand, better interpret the neural response to the presented experimental stimuli. In a previous study Hölle et al. (2021), we could not say whether a missed response to a target sound was due to a lapse of attention or due to the acoustic environment. Recording the participants soundscape can uncover energetic masking of the stimuli, thereby extending our previous work. On the other hand, we can compute the neural response to naturally occurring sounds.

We are specifically interested in discrete, transient sounds - a door closing, a phone starting to ring, a car honking - from which we can expect that they elicit clear auditory event-related potentials (ERPs). Auditory ERPs are categorized into components. The N1 component indicates the degree to which a sound has been processed (May & Tiitinen, 2010). In presenting paired click sounds, we can also quantify the amplitude attenuation from the first to the second sound, which can be interpreted as filtering out irrelevant information (Bak et al., 2017; Jones, Hills, Dick, Jones, & Bright, 2016; Major et al., 2020). The P3 component is associated with attention to stimuli, i.e., sounds that are relevant for a person (Polich, 2007). Taken together, ERP components provide insights into the topdown and bottom-up processes. Components such as N1 and P3 can be recorded with ear-EEG (Bleichner, Mirkovic, & Debener, 2016; Debener, Emkes, De Vos, & Bleichner, 2015; Denk et al., 2018; Holtze, Rosenkranz, Jaeger, Debener, & Mirkovic, 2022; Meiser & Bleichner, 2022; Meiser, Tadel, Debener, & Bleichner, 2020; Mirkovic, Bleichner, De Vos, & Debener, 2016), even beyond the lab(Hölle et al., 2021; Somon et al., 2022).

In this study, we recorded mobile ear-EEG for four hours using a mobile smartphone-based ear-EEG setup. For one hour, participants completed four conditions in the lab where they listened to paired-click sounds with different instructions. These conditions serve as a reference to compare the beyond-the-lab condition to. In the beyond-the-lab condition, participants follow their daily activities in an office for three hours while they are sporadically presented with blocks of paired-clicks. They receive no further instructions. Additionally, all environmental sounds were recorded. We want to investigate to which of the reference recordings the beyond the lab recording is most similar to and how environmental sounds were processed.

## 2 Methods

### 2.1 Participants

Twenty-four healthy undergraduates (13 female, 11 male), age ranged from 21 to 30 (*M* = 24.71, *SD* = 2.49), were recruited via the university’s online blackboard or by word-of-mouth. We made no prior calculations regarding sample size in this exploratory study. Participants were compensated with 10€ per hour. Due to technical reasons, the data from nine participants had to be excluded completely from further analyses (see discussion section for a detailed discussion). From five other participants, only parts of the data could be used: one participant lacked one condition and four participants lacked acoustic information. Prior to participation, all participants provided informed consent. The study protocol was approved by the ethics committee of the University of Oldenburg.

### 2.2 Paradigm

We performed auditory experiments under controlled conditions in the lab for one hour and beyond the lab in an office situation for three hours. Specifically, the experiment consisted of four conditions in the lab (task-free, counting, reading, listening) and the beyond-the-lab (BTL) condition in the office (see Figure 1 for an illustration).

**Figure 1:**
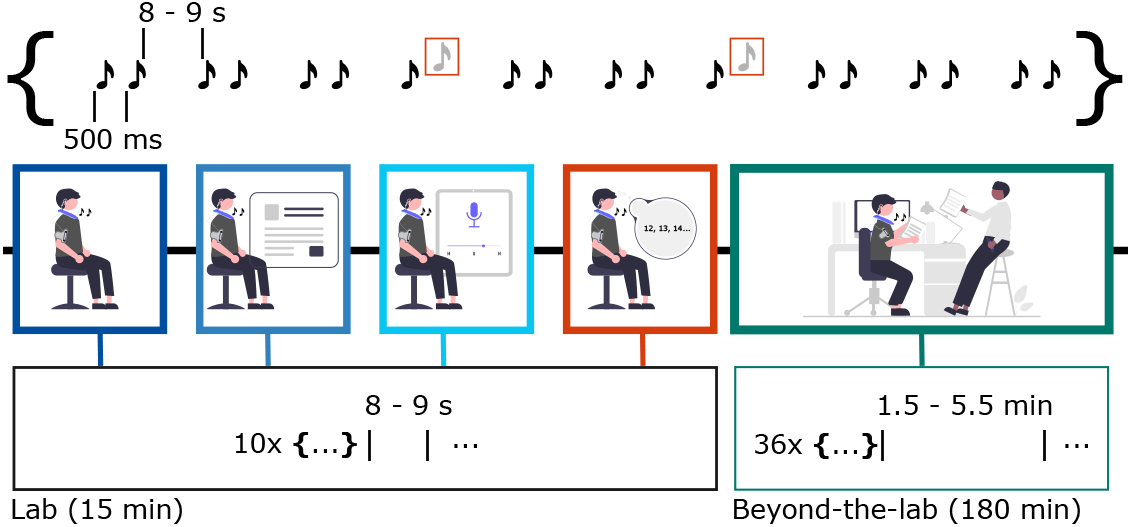
Overview of the paradigm. Participants completed four conditions in the lab (task-free, counting, reading, listening) and one condition in the office (beyond-the-lab) while click-pairs were presented. In the lab condition, they were presented with to a total of 100 clicks (80 standard, 20 deviant) in quick succession. In the beyond-the-lab condition, they were presented with a total of 360 clicks (80 standard, 20 deviant) that occurred in sporadic sequences of ten clicks. Note that deviant sounds were only relevant for the counting condition and were discarded in all analyses. The order of the four lab conditions and the beyond-the-lab condition was counterbalanced.

In all of these conditions, we used a paired-click paradigm (Gjini et al., 2011). Each click pair consisted of two click tones 500 ms apart. In 80% of click pairs (standard), the first and the second click were identical (1000 Hz, 4 ms duration, 1 ms onset and offset ramps). In 20% of the click pairs (deviant) the second tone differed in pitch and duration (2500Hz, 8 ms duration, 2 ms onset and offset ramps). Note, in all our analyses we only analyzed the standard clicks, i.e., the trials in which both clicks were acoustically identical.

In each of the four lab conditions, 100 click-pairs were presented at an intertrial interval that varied from 8 to 9 s (*minimal standard random number generator*), lasting for a total of 15 minutes. The order of standard (80% of all trials) and deviant pairs (20% of all trials) was randomized with the restriction that no more than two deviant pairs occurred successively. In the BTL condition, in total 360 click-pairs were presented in 36 blocks, henceforth referred to as sequences. Each sequence consisted of 10 click-pairs (8 standard, 2 deviant) that were presented every 8 to 9 s in randomized order. The time between sequences varied from 1.5 minutes to 5.5 minutes. In total, the BTL condition took approximately 180 minutes.

The lab conditions only differed in the instructions to the participant; in other words, the sound stimulus in all conditions were identical, but participants had to behave differently towards them. In the task-free condition, participants were instructed to look at a fixation cross at eye level while clicks were presented. In the reading condition, they read a German newspaper article about introverted employees (*Leiser, bitte!*, Die Zeit, Zeitverlag Gerd Bucerius GmbH & Co. KG, Hamburg, Germany). In the listening condition, they listened to a German audio article about marriage (*Warum Heiraten?*, Die Zeit, Zeitverlag Gerd Bucerius GmbH & Co. KG, Hamburg, Germany, presented at ~ 60 dB on Sirocco S30 loudspeakers, Cambridge Audio, London, United Kingdom). In the counting condition, they had to count the deviant pairs. All lab conditions took place in a quiet room.

In the BTL condition, participants did not receive any instructions on how to behave towards the clicks and could perform a self-chosen task in an office room where no other person was present (due to Covid-restrictions). Participants performed activities such as reading, studying, using the smartphone, watching movies, or working on the laptop. Their only restrictions were to not leave the room (except for toilet breaks) and to not use headphones, as our microphones would not record the auditory stimulation by headphones. This condition also included a brief trip along a street to the crowded cafeteria with the experimenter around lunch time. The experiment continued running during lunch. Refer to Figure 2 for an illustration of the soundscape and movement over time in this condition. The time courses of each participant can be found in supplementary Figures 1–11.

**Figure 2:**
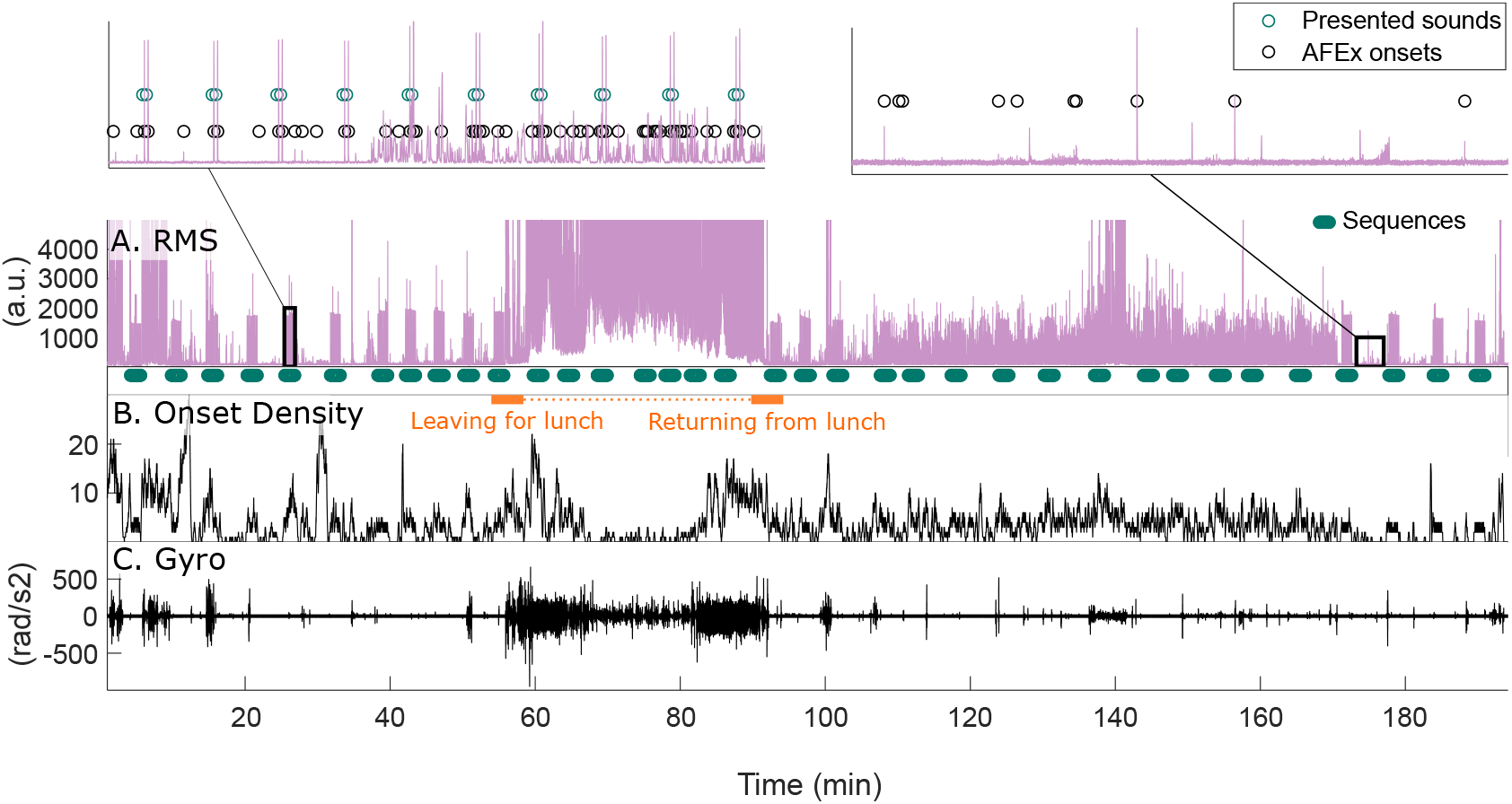
Time courses of RMS, sound onsets (as detected by the AFEx app), and movement of one participant in the BTL condition. **A.** RMS time course (only one of the two stereo streams is shown). Below, the occurrence of sequences (multiple presented sounds) is shown. The zoomed RMS segment on the left shows presented sounds (paired-clicks) and AFEx onsets. It is visible that presented sounds also generate AFEx onsets. On the right, a segment where no sounds were presented, but AFEx onsets were detected, is shown. **B.** Time course of onset density. It was calculated by a moving window of 10 seconds in which all onsets were summed up. The interval in the cafeteria is indicated by the orange bars. During this interval, the number of detected AFEx onsets decreases due to the higher level of background noise (no sound events stand out) **C.** Time course of the gyroscope data (only one axis is shown). Periods of more movement, for instance, on the way to and from the cafeteria, are indicated by the higher deflections.

The order of conditions was counterbalanced with the following restrictions: first, participants either started or finished with the BTL condition. Second, the lab conditions always started with the task-free condition. These restrictions resulted in 12 unique permutations. The order of the lab conditions ensured that the task-free condition was not influenced by the instructions of the other conditions. If participants had started with the counting condition, they could have counted still in the task-free condition. Note also that in the BTL condition, the lunch time was kept approximately constant in each counterbalance order (12 PM if they started with the BTL condition, 1 PM if they started with the lab conditions).

### 2.3 Apparatus

Participants were equipped with a neckspeaker with an integrated amplifier (nEEGlace), two cEEGrids, a smartphone worn on the arm, and ear-microphones. The setup is illustrated in Figure 3.

**Figure 3:**
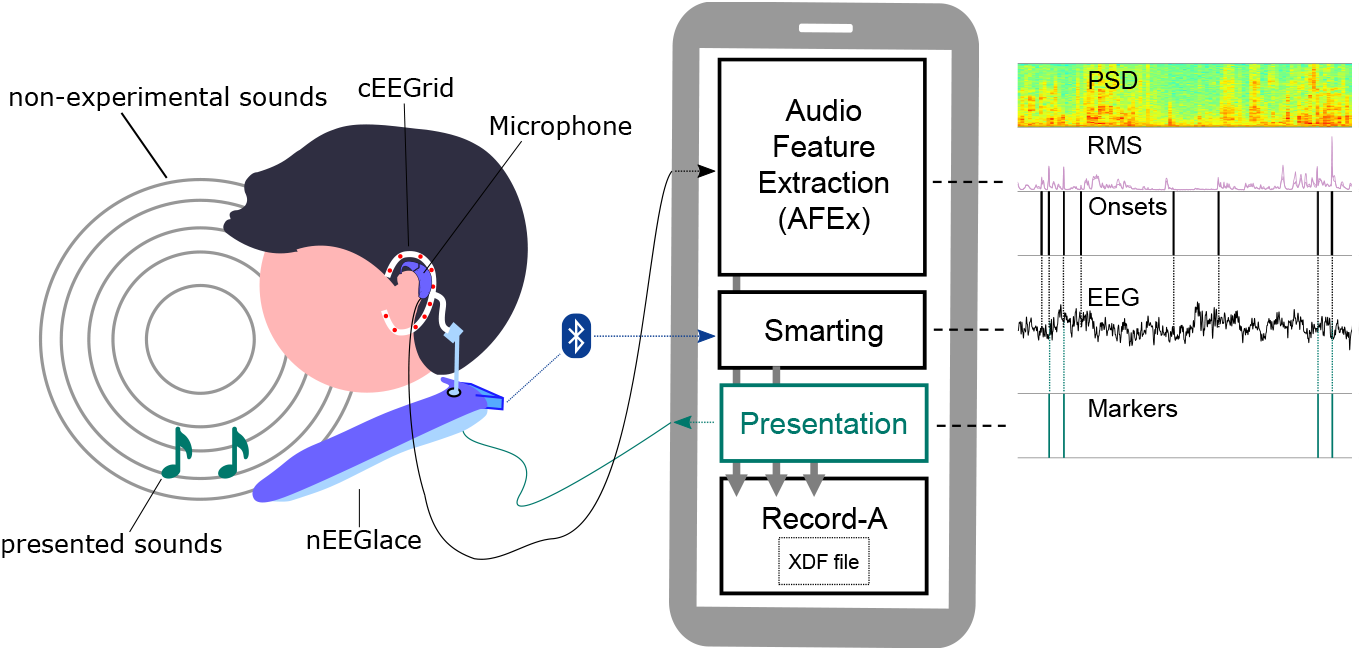
Illustration of the experimental setup. The participant was equipped with cEEGrids, the nEEGlace, ear-microphones, and a smartphone. EEG was recorded with cEEGrids and transmitted via Bluetooth to the smartphone with the Smarting app. Experimental sounds were presented via the nEEGlace connected to the smartphone via audio cable and stimulus presentation was controlled by the Presentation app. Non-experimental sounds were recorded via ear-microphones connected to the smartphone and acoustic features (PSD, RMS, onsets) were extracted by the AFEx app. Resulting datastreams are shown to the right of the smartphone. Both the markers from presented sounds and from AFEx onsets can be used to analyse the EEG data. All datastreams were recorded by the Record-a app into a single xdf-file.

The nEEGlace (pronounced necklace) is a custom-made EEG system consisting of a neckspeaker with an integrated research-grade 24-channel Smarting (mBrainTrain, Belgrade, Serbia) amplifier with 24 bit resolution and a customized connector for cEEGrids (for details, see Bleichner & Emkes, 2020). The amplifier has an integrated 3D gyroscope. Around each ear of the participant, we placed one cEEGrid (tMSI, Oldenzaal, The Netherlands) with ten Ag/AgCl electrodes (DRL: R4a, Reference: R4b). The EEG signal was sampled with 250 Hz and the data was transmitted via Bluetooth to the Smarting Android app (Version 1.6.0) on the smartphone.

The smartphone (Google Pixel 3a, OS: Android 10) was used for data recording and for stimulus control. It was connected to the nEEGlace via an audio cable. Click sounds were presented with the Presentation Android app (Version 3.0.0, experiment programmed with NBS Presentation Version 22.0; Neurobehavioral Systems, Inc., Berkeley, CA, USA) and presented by the nEEGlace (LAFmax ~ 77 dB). Consequently, as opposed to wearing headphones, the participant could still hear all environmental sounds. The Presentation app generated markers for the click sounds.

To record sound at ear-level, we used customized ear-microphones (two omnidirectional EK-23024 microphones, Knowles Electronics, Illinois, USA, placed into behind-the-ear shells). The ear-microphones were connected to the phone via the USB-C port with a sound adapter (Andrea Pure Audio USB-MA, Andrea Electronics, Bohemia, USA) and a micro-USB to USB-C adapter (Hama, Monheim, Germany). For sound recording, we used the open-source Android app AFEx(Hölle et al., 2022) that records environmental sounds as privacy-protecting acoustic features. The app records sound onsets, henceforth referred to as AFEx onsets, average signal power (RMS), and power spectral density (PSD).

All three apps (Smarting, Presentation, AFEx) generated datastreams by using the Lab Streaming Layer (LSL). All datastreams were recorded by the Record-a app (Blum, Hölle, Bleichner, & Debener, 2021) into a single xdf-file.

### 2.4 Procedure

All participants arrived in the lab at approximately 10 AM with their hair washed. The skin around their ear was cleaned with abrasive gel and alcohol swabs. A small strip of Leukosilk medical tape (BSN medical GmbH, Hamburg, Germany) was placed behind their ear to protect the skin and increase comfort. Small drops of electrolyte gel (Abralyt HiCl, EasycapGmbH, Germany) were applied to each cEEGrid electrode and the cEEGrids were placed around the ear with double-sided adhesive stickers. Impedance of most electrodes was below 10 kOhm at the start of the recording or approached this threshold during the recording. Participants were equipped with the nEEGlace around their neck and cEEGrids were connected to the amplifier. The smartphone was connected to the nEEGlace via audio cable and the ear-microphones were attached to the smartphone via the USB-C port. The microphones were placed on top of their ears (similar to hearing aids) and the smartphone was stored in an arm-pouch that was attached to their arm.

Before the experiment started, participants performed a short three-minute oddball experiment in which they had to count rare, deviant beep sounds (not the click sounds used in the experiment). The data from this oddball was analysed right afterwards (not reported) to make sure that the setup was functional and the data quality was sufficient to proceed with the experiment. If the typical ERP responses (N1 and P3 to deviant beeps) were visually identified and cEEGrid channels showed good data quality, the experiment was started.

Participants either started with the conditions in the lab or with the BTL condition (see paradigm). Each condition started with a one-minute calibration phase where participants were instructed to look at a fixation cross at eye level. Before each condition, the experimenter explained the task and clarified questions. After both the reading and listening condition, participants had to answer seven content-related multiple choice questions (not reported).

After all conditions were recorded, the setup was removed, participants cleaned up residual gel around their ears with paper tissues. Lastly, they completed the Noise Sensitivity Questionnaire (NoiSeQ; Sandrock, Schütte, & Griefahn, 2007; Schütte, Marks, Wenning, & Griefahn, 2007), a 35-item questionnaire that measures noise sensitivity on a Likert-type scale coded from 0 (strongly agree) to 3 (strongly disagree). The global score was calculated by taking the average of all items.

### 2.5 Data Analysis

For the offline data analysis, we used Matlab (Version 9.6.0.1335978; The Mathworks Inc., Natick, MA, USA), the toolbox EEGLAB (Version v2019.0; Delorme & Makeig, 2004) including the *cEEGridplugin* (Version 0.9, https://gitlab.com/mgbleichner/ceegridplugin) and custom scripts.

#### 2.5.1 Preprocessing

The EEG data was low-pass filtered at 25 Hz (Order 133) and high-pass filtered at 0.1 Hz (Order 8251). The filters were used as implemented in EEGLAB (*pop_eegfiltnew*). Identical to Hölle et al. (2021), we performed artifact subspace reconstruction (ASR) as implemented in *clean_rawdata* (Version 1.0; parameters: flatline criterion = 60, high-pass = [0.25 0.75], channel criterion = off, line noise criterion = off, burst criterion = 20, window criterion = off) to clean artifacts. Rejected channels based on the flatline criterion (*M* = 1, *SD* = 1.9) were spherically interpolated.

**Events** The datasets contains events for AFEx onsets (sound onsets generated by the AFEx app) and for presented sounds (clicks). Note, the sounds presented with the nEEGlace were also captured by the microphone and hence are present both as presentation markers as well as AFEx onsets. To account for the known buffering lag of the AFEx app, we shifted all AFEx onsets by −248 ms (i.e., 62 samples, see Hölle et al., 2022). Additionally, for all events in the dataset (presented sounds and AFEX onsets), we computed an average RMS value consisting of the mean of of 62.5ms (5 samples) after the event. This average RMS value was used as a measure of the background noise during each event.

#### 2.5.2 Analysis of presented sounds

Epochs were extracted from −0.1 to 1 s (baseline −0.1 to 0 s) relative to the first click of the standard click-pair (80 epochs for each lab condition and 288 epochs for BTL) and then averaged. Note that we only analyzed the standard sounds; therefore, from the 100 lab and 360 BTL trials only 80% are analysed.

The BTL condition contained more movement of the participant compared to the lab conditions and a changing soundscape, decreasing the signal-to-noise (SNR) ratio. We therefore also computed a corrected average of the BTL condition where we removed epochs with extensive background noise and movement. Movement per epoch was calculated by taking the average over the epoch of the RMS of all three gyroscope axes. We then calculated a RMS and movement threshold based on the 85th quantile over all epochs and participants. Epochs exceeding this threshold were removed (*M* = 25.76%, *SD* = 11.35%).

**Sequences** In the BTL condition, clicks were presented in sequences consisting of eight standard click pairs (see paradigm). For some analyses (see classification and statistical analysis), we investigated the data at the sequence level to counteract the limited SNR of single trial ear-EEG data (Meiser & Bleichner, 2022; Meiser et al., 2020). Therefore, for all sequences, we averaged the eight click epochs to obtain a local ERP. Before epochs from a sequence were averaged, we rejected bad epochs (M = 8.30%, *SD* = 1.00%) based on joint probability with a threshold of two standard deviations. Additionally, we also computed the average RMS and movement for each sequence (as opposed to for each single epoch).

#### 2.5.3 Generalized eigendecomposition

To include information of all recorded channels, we used generalized eigende-composition (GED, Cohen, 2022) to obtain a spatial filter that maximises the amplitude of the N1 compared to baseline. To compute the GED, we concatenated the epochs from all participants from the lab conditions, as these controlled, motion-free conditions provide the best data quality. To maximize the amplitude of the N1, we contrasted the time window of the the N1 component (100 ms to 160 ms) to the baseline period (−100 ms to 0 ms) and then extracted the spatial filter that explained the most variance. For all ensuing analyses and all figures, we used this GED channel (see Figure 4).

**Figure 4:**
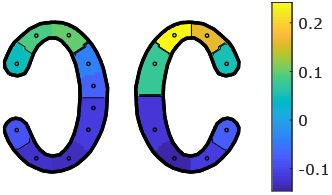
Virtual channel based on generalized eigendecomposition (GED). This GED channel is used for all analyses.

#### 2.5.4 Analysis of AFEx onsets

We extracted epochs from −0.4 to 0.5 s (baseline −0.1 to 0 s) relative to each AFEx onset in the BTL condition. Note that we chose a different time window than for the presented sounds here for two reasons: first, we expect the neural response (P1-N1-P2-P3) to an AFEx onset to fade after approximately 500ms. Second, we wanted to exclude the overlap of neural responses due to AFEx onsets that are close to each other in time (see below). The total number of AFEx onsets ranged from 2975 to 10685 over all datasets (*M* = 5485.3, *SD* = 2027.5). On average, an onset was detected every 2.17s (SD = 4.42s).

Importantly, the presented click sounds also generated AFEx onsets (see Figure 2). As we were interested in natural, non-experimental sound onsets, we identified AFEx onsets generated by presented sounds and removed them for the analysis of non-experimental sounds. They were identified by finding the closest AFEx onset around the respective click marker. However, these AFEx onsets are still informative to validate the detection of onsets.

After removing epochs of AFEx onsets generated by presented sounds, we discarded those epochs that contain more than one AFEx onset due to the overlap (M = 33.45%, *SD* = 0.08%). Remaining bad epochs were rejected (M = 5.11%, *SD* = 1.11%) based on joint probability with a threshold of two standard deviations. Lastly, to exclude remaining noisy epochs, we rejected epochs exceeding a movement threshold (M = 9.22%, *SD* = 1.10%) based on the 85th quantile of the mean movement over all epochs per participant.

As we expected clear ERPs (showing P1-N1-P2 complexes) from the presented sounds, we were also interested in non-experimental sounds that were similar in PSD and RMS to the presented sounds. To determine similarity, we used the following procedure: First, the sound onset had at least the same or an 2000 a.u. higher RMS value than the presented sounds. Second, the correlation of the sound onset and a PSD template consisting of nine samples (1.13 s) relative to the presented sound was higher than the computed threshold. To gain the threshold, the PSD template was correlated with the PSD representation of every sound onset for each participant. The correlation threshold was then based on the 80^th^ quantile of this correlation distribution. With this procedure, we identified, on average, 321.73 (SD = 251.02) epochs. We then rejected bad epochs based on joint probability (threshold: 2 SDs; *M* = 7.85%, *SD* = 1.80%) and movement (exceeding the 85th quantile of the mean movement over all epoch; *M* = 13.83%, *SD* = 0.32%).

#### 2.5.5 Noise estimation

To obtain an estimation of noise in the EEG signal in the BTL condition, we computed an amplitude threshold based on random events that were inserted into the dataset. For each participant, for 1000 permutations we inserted events at random time points. We used these markers for epoching and calculating the grand average ERP. After rejecting epochs based on joint probability (threshold: 2 SDs), we averaged over all permutations per participants and over all participants and time points. The amplitude threshold consists of this mean plus-or-minus 1.96 times the standard deviation.

#### 2.5.6 Classification

To understand how sequences in the BTL condition were processed with regards to the lab reference conditions, we used a quadratic discriminant classifier. For each participant, we trained a classifier on the data from the lab and then tested it on the data from the BTL condition.

The classifier was trained on peak amplitude ERP features from the lab conditions. To match the SNR of the sequences in the BTL condition (see sequences), we used a moving average over eight trials; hence, each new trial consisted of itself and seven subsequent trials averaged. We then extracted peak amplitudes for the P1, N1, P2, and P3 for both the first and the second click, resulting in eight features that were extracted with the following method: first, for each participant, we empirically determined the peak latency based on the condition average by searching in a specified time window after a click (P1:40-90 ms, N1: 100-160 ms, P2: 180-260 ms, P3: 300-400 ms). Second, for each trial, we computed a 20ms average around the identified peak latency.

The same features were extracted from each sequence in the BTL condition with the identical procedure.

We combined the three lab conditions that did not require to focus on the clicks (task-free, reading, and listening, all correlate with 0.74-0.90), hereafter referred to as passive condition. The counting condition, in contrast, is referred to as active condition. Consequently, the classifier classified sequences from the BTL condition as either belonging to the passive condition or the active condition.

To train the classifier with the same number of trials for both the active and the passive condition and to better interpret and validate the classifier, we randomly subsampled trials from the passive condition for each participant. Hence, we trained 100 classifiers with a different subsample and then tested each of these 100 classifiers on the BTL sequences. When the majority (more than 50%) of all classifications labeled a sequence as belonging to the passive group, for instance, it was accepted as passive.

Ten-fold cross-validation was used to evaluate the classifier performance. The mean prediction error over permutations ranged from approximately 2% to 20% (M = 11.27%, *SD* = 5.17%).

We also performed a quadratic discriminant classification trained on all four conditions. The results of this classifier is shown in supplementary Figure 12.

#### 2.5.7 Statistical analysis

We performed the statistical analyses with Matlab and RStudio (Version: 2022.02.1; RStudio, PBC, Boston, MA, USA; R-Version: 4.1.3). For all tests, an alpha level of 0.05 was used (Bonferroni-corrected in case of multiple comparisons). We analysed the EEG data from the BTL using linear mixed models (LMM) with the R-packages *lme4* (Bates, Mächler, Bolker, & Walker, 2015) and *lmerTest* (Kuznetsova, Brockhoff, & Christensen, 2017). Reported p-values were determined by using the Satterthwaite approximation (Volpert-Esmond, Page-Gould, & Bartholow, 2021).

In the BTL condition, we were specifically interested in masking effects of the presented sounds due to background noise (cf. Hölle et al., 2021). For this purpose, we first identified sequences with high background noise and low background noise by performing a median split of the RMS per sequence over all participants. We then build an LMM for each ERP component (P1,N1,P2,P3) for the first tone with the fixed factor *Background Noise* and a random intercept for *participant*. Due to non-convergence of the model for the P2 and P3 component, we simplified the model by removing the random intercept (simple linear model).

To statistically analyze an amplitude attenuation from the first click to the second one in a click pair, we quantified the N1 for each participant in each condition by computing a 20 ms average around the peak latency. We identified the peak latency based on the condition grand average in the time window from 100 to 160 ms after each click. For each condition, we performed a paired t-test with the amplitudes of click 1 and those of click 2.

## 3 Results

### 3.1 NoiSeQ

The mean score of the NoiSeQ was 1.5 (*SD* = 0.25). Due to the homogeneous scoring of the sample, we do not distinguish between high and low scorers in any of our analyses.

### 3.2 EEG

Figure 5A-E shows the grand average ERP for all conditions. We observed auditory evoked potentials (P1,N1,P2) in all conditions. As expected, the morphology and amplitudes are different across conditions. Generally, the overall amplitudes decrease over the passive conditions (task-free, reading, listening). In the task-free and reading condition, a significant N1 amplitude attenuation from the first to the second click can be observed (task free: *t*(14) = −3.56, p<0.01; reading: *t*(14) = −4.00, *p*<0.01). We found no significant attenuation in the listening, oddball, and BTL condition (all p>0.15). Specifically in the counting condition, the average amplitude increased to −19.57μV in the second tone compared to −12.50μV in the first tone, which can be explained by the potentially deviant second click that had to be counted by participants. Visually, the P2 component to the first tone retains a similar amplitude over all lab conditions. In the BTL condition (Figure 5E), we show the average of all trials and the average with trials corrected for movement and background noise in overlap. The small differences between both ERPs indicates that the noise caused by movement and background noise is averaged out even when not explicitly correcting for it.

**Figure 5:**
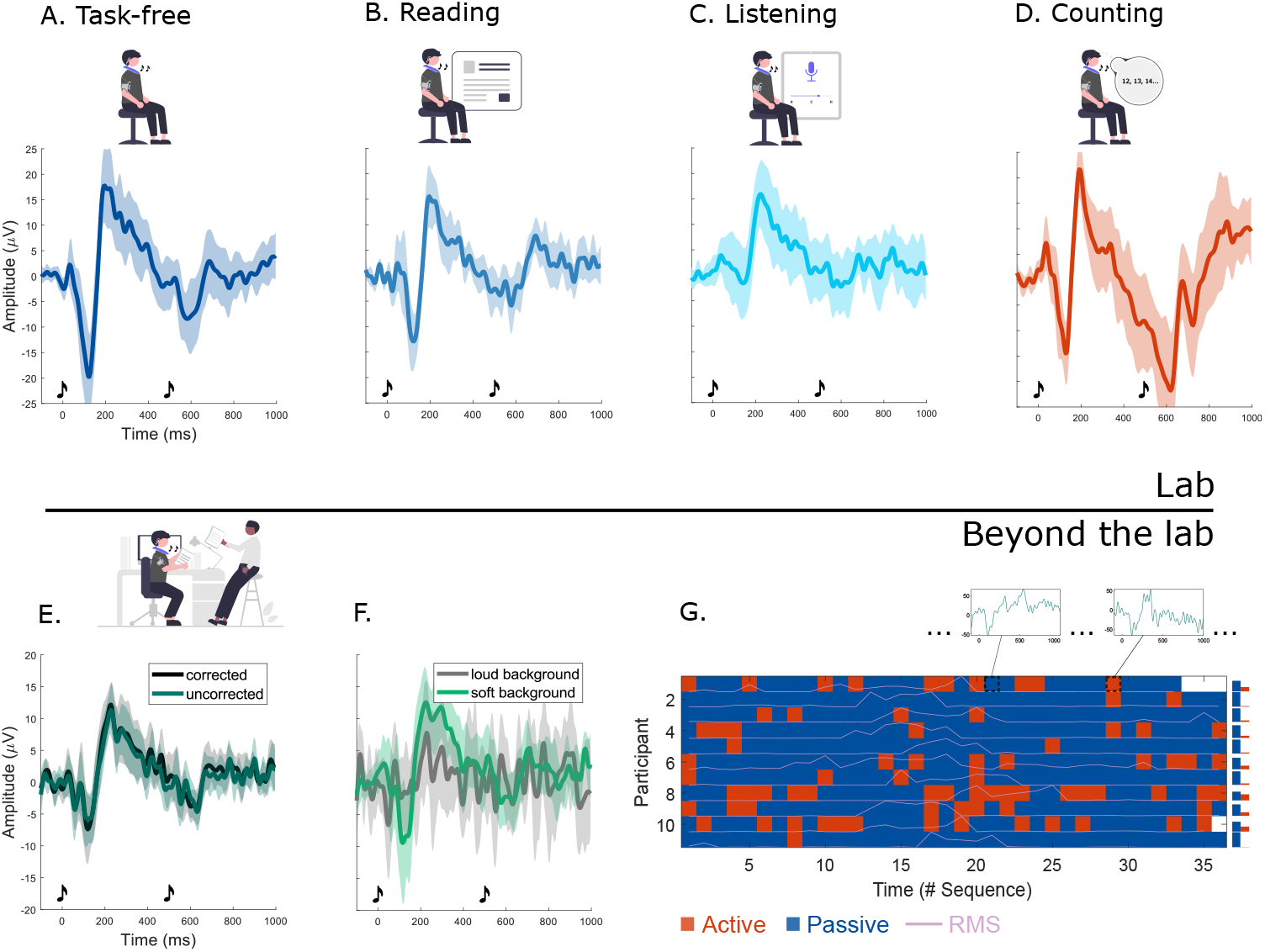
Grand average ERPs of all conditions. Shaded areas indicate the 95% confidence interval. **A.** Listening to clicks without instructions (n=15). **B.** Reading a newspaper article while clicks are presented (n=15). **C.** Listening to an audio article while clicks are presented (n=15). **D.** Counting deviant click pairs (n=14). **E.** Performing a self-chosen activity in an office room while clicks are presented (n=15). Green: All trials; Black: Trials free from high levels of movement or background noise **F.** ERPs from the beyond-the-lab condition split in trials with high and low background noise (n=11). **G.** Results of the classification (n=11) of each sequence based on the lab reference conditions (passive: task-free, listening, reading; active: counting). Two exemplary sequences are visualized above. For each participant, the RMS trace (normalized to the maximum) is shown in overlap. Note that two participants have missing sequences at the end due to battery failure. The bars on the right visualize the percentage of each mode per participant

Furthermore, Figure 5F illustrates the grand average of sequences with high background noise compared to sequences with low background noise. The mean amplitude of the N1 during low background noise was −12.95*μ*V (*SE* = 3.15) with a mean increase of 7.17*μ*V (SE = 3.58) for high background noise (*p*<0.05). The mean amplitude of the P3 (M = −11.57 *SE* = 2.32) during low background noise showed a decrease of −6.81*μ*V (SE = 3.28) for high background noise (*p*<0.05). No significant difference emerged from the P1 and P2 component. In other words, both the N1 and the P3 component were more pronounced during low vs high background noise, indicating a masking effect. The full results of the models can be found in supplementary Table 1–4.

The results of the classification are shown in Figure 5G. We classified each sequence based on our lab reference conditions for each participant. The figure illustrates that participants switched between the active and the passive condition, but most sequences were classified as passive. For each participant, we also show the normalized RMS timecourse in this plot.

In Figure 6, we show the results of the analysis of the AFEx onsets from the BTL condition. In Figure 6A, we show the grand average of all AFEx onsets. Note that no trials were rejected in this plot. Figure 6B illustrates the AFEx onsets that were generated by the presented click sounds (see Analysis of AFEx onsets). Since we know exactly when we presented click sounds due to the respective markers, we can use the AFEx onsets generated by these sounds to validate the onset detection. On average, AFEx detected an onset event for 78.74% (SD = 10.04%) of all presented sounds. The computed average based on these AFEx onsets correlates with the grand average of the BTL condition (shown in overlap, also see Figure 5E) with 0.95. Due to this high correlation of the grand average of presented sounds and the grand average of AFEx onsets generated by these presented sounds, we can safely assume that the timing of the AFEx onsets is accurate. Figure 6C shows the average of AFEx onsets that were acoustically matched to the presented sounds based on RMS and PSD. Figure 6D depicts the 200 loudest AFEx onsets based on RMS per participant. Neither Figure 6C nor D show auditory evoked potentials. Below the ERPs in 6B-D, we also show the grand average RMS in the identical time window.

**Figure 6:**
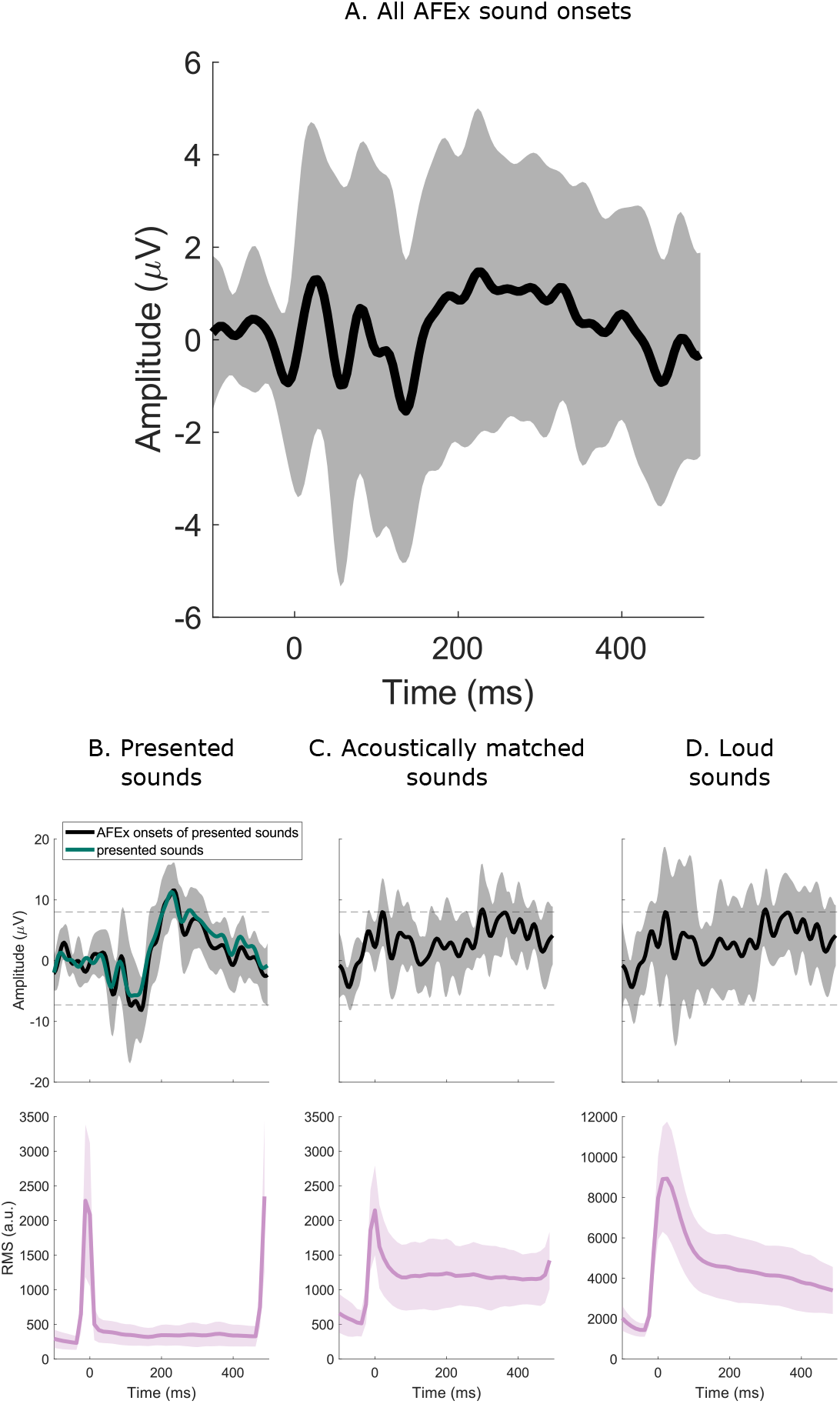
Grand average ERPs of AFEx onsets generated by presented (A) and naturally occurring sounds (B,C). The shaded areas indicate the 95% confidence interval. **A.** All AFEx onsets **B.** Black: AFEx onsets generated by presented sounds (click stimuli); Green: ERPs based on marker of presented sounds. **C.** AFEx onsets similar to presented sounds matched on loudness and PSD **D.** 200 loudest (based on RMS) AFEx onsets per participant. The dotted black lines represent a noise estimate. Below the ERPs, the corresponding grand average RMS is shown.

## 4 Discussion

The goal of this study was to investigate how individuals process sounds in everyday life. Here, we used smartphone-based mobile ear-EEG to monitor sound processing in the lab and while participants worked unsupervised in the office. We studied ERPs in response to controlled click sounds as well as in response to naturally occurring sounds. Thereby we could compare the beyond-the-lab results to reference recordings of the same participant using the same hardware obtained under controlled lab conditions. During the lab recordings, we instructed the participant to behave differently towards the sound source. Beyond the lab, we did not provide the participants with any instructions. In the following, we first discuss our results regarding the presented sounds followed by the analysis of the natural sounds.

The lab conditions that all used the same stimuli but different instructions showed auditory ERPs with different morphologies and amplitudes, which are highly consistent with the literature (cf. Gjini et al., 2011; Major et al., 2020; May & Tiitinen, 2010; Rosburg, Trautner, Elger, & Kurthen, 2009; Shepherd et al., 2016). Across all conditions, we observed an N1 and P2 in response to the first click sound. Generally, the amplitude of the N1 depends on the task, whereas the P2 appears to be stable across conditions (cf. Gjini et al., 2011; Major et al., 2020; Shepherd et al., 2016). In the task-free and reading condition, the N1 to the second tone shows an amplitude attenuation compared to the first tone (cf. Gjini et al., 2011; Major et al., 2020; May & Tiitinen, 2010). In contrast, in the counting condition, in which the second tone could be relevant, no amplitude attenuation was found; in fact, the amplitude increased from the first to the second tone (cf. Rosburg et al., 2009). In the listening condition, this amplitude attenuation is not apparent and amplitudes are generally smaller compared to the other conditions. This finding is consistent with Major et al. (2020) who also found smaller amplitudes and no N1 attenuation in their listening condition compared to other conditions. In sum, the cEEGrid can measure differences in ERPs generated by different instructions.

Importantly, we also find comparable ERPs in the BTL condition in response to the presented sounds. While the same controlled stimuli were used as in the lab condition, the acoustic environment and the behavior of the participant were uncontrolled, and the participant had no explicit instruction towards the presented stimuli. As in the lab conditions, we see an N1 and P2 for the first tone, and a reduction in amplitude of the N1 for the second tone, approaching significant levels. Hence, we can capture the neural response to specific sounds in freely behaving participants without explicit instructions towards the stimulus. These results therefore extend our previous study (Hölle et al., 2021) where we recorded mobile long-term EEG in the office while participants performed an active oddball task, i.e., a task in which they had to attend to all presented sounds explicitly. Furthermore, in the current study, we used the AFEx app to record the background noise level and could therefore explore the effects of masking. As expected, the neural response for our presented sounds was smaller during high background noise compared to low background noise. Hence, in noisy soundscapes the clicks were potentially not perceivable. However, we do not know whether these masking effects are due to energetic masking (i.e., it was too loud to hear the tones) or due to informational masking (i.e., participants paid attention to the background noise, Arbogast, Mason, & Kidd, 2002; Brungart, 2001). Taken together, we can conclude that we can measure the neural response to presented sounds in everyday situations if the sounds stand out against the background.

Auditory functioning varies during the course of day (Basinou, sub Park, Cederroth, & Canlon, 2017). We thus also explored the sound processing of an individual resolved over time. Does the sound processing vary over time and situations? In an exploratory analysis, we therefore used a classification approach. The results suggest that participants’ sound processing as indexed by their ERPs resemble the passive listening conditions most of the time, but occasionally switches into an ‘‘active” processing mode. Admittedly, these results are provisional and merely serve here as an illustration of how to approach individual sound processing in everyday life. The presented analysis suffers from two major limitations. First, we have no ground truth to which we can compare the results to, that is, we do not know, how people perceived these sounds. Second, we did not track the participants’ behavior during the BTL condition. We do not precisely know when they read, listened, or watched, which could have added valuable information to the BTL condition.

In future studies, we advise to gain a better insight into a person’s behavior to aid the interpretation of results. There are several possibilities to do so: first, one could observe participants, they could be filmed (cf. Scanlon et al., 2019), or sound snippets could be randomly recorded via app (Mehl, 2017) to code their activity - but both approaches will invade the participants’ privacy. Second, movement as recorded by a gyroscope or additional (smartphone) sensors could be used for activity recognition (Ramasamy Ramamurthy & Roy, 2018). Third, following an experience sampling approach (Holube, von Gablenz, & Bitzer, 2020; Trull & Ebner-Priemer, 2013), participants could self-report their current activity on the smartphone at several occasions over time. All these approaches come at the potential cost that they alter natural behaviour. When the behaviour of the participant is recorded by video, microphone, motion tracking, or self-report participants might change their behavior, e.g., to conform to some perceived socially desirable behavior. Overall, a balance between necessary information for the research question, undisturbed behaviour, and privacy concerns is required.

In sum, we have successfully extended our previous work: We found different task-dependent ERPs to click stimuli in all lab conditions measured by ear-EEG. Beyond-the-lab, we found a comparable ERP response without any task instructions and masking effects dependent on background noise level.

Obtaining high-quality datasets beyond the lab remains challenging. Even though we ensured to have a sufficiently high recording quality in the beginning of the recordings, we still lost more than one third of the recordings, due to poor data quality and malfunctioning of hardware. There are several factors that might cause poor data. Mechanical problems can interrupt the signal transmission from head to the amplifier, such as connections becoming loose or bends in the cables. Furthermore, the electrode skin interface may decrease due to hair or sweating. Lastly, participants may have changed the setup by mistake. We aim to improve these aspects with revised setups in the future by reducing the number of wires and by developing new cEEGrids. With the current setup, a high level of attention is needed during all steps of experimenting and especially during preparation. To counteract the loss of datasets in future studies, we have produced a video tutorial (Hölle & Bleichner, 2022) on how to acquire high-quality ear-EEG data recorded BTL that provides best practice recommendation and increases standardization across different experimenters.

With the AFEx app, we detected onsets, recorded spectral information and RMS. We have thus extended the results of Höolle et al. (2022) by showing that we can track sound events in real life. By using the presented stimuli, we could validate the onset detector in real life. When using the AFEx onsets instead of the presentation markers, we could obtain comparable ERP responses, underlining the general functionality of the system. Interestingly, we did not find a clear neural response to the natural, non-experimental sounds. This finding was unexpected as we have shown in a previous study that we can obtain clear AFEx onset based ERPs in response to natural sounds in a controlled lab condition (Hölle et al., 2022). There are several possible explanations for this discrepancy. First, in Höolle et al. (2022) participants listened passively to a presented soundscape in the lab; in the current BTL condition, participants were not restricted in their behaviour and might have produced some of the sounds that were detected as onsets by themselves. It is well established that self-generated sounds lead to a reduced neural response (Cao, Thut, & Gross, 2017; Sanmiguel, Todd, & Schröger, 2013; Schneider & Mooney, 2018). From the limited information provided by the acoustic features (sound onsets, RMS, PSD), we could not determine whether a sound was self-generated by the participant or generated by the environment. Second, many of the presented sounds could have been predictable (e.g., observing another person making a sound). This is different to both the presentation of an everyday life scene in the lab and to the click sounds presented in this study that occurred without cues or context. The click task actually provides an indication how strong these expectancy effects can be, as the N1 to the second tone, i.e., the one that can be expected after the first tone is heard, is greatly reduced. Lastly, everyday life sounds are not only very different from sounds that are experimentally presented, but also have larger variability. The presented sounds have sharp onsets and offsets made to elicit clear ERP; in natural sounds, such features are seldom seen (see RMS of Figure 6).

To understand sound processing in everyday life we need more information about the context and behaviour. One possible approach is to use auditory scene analysis (Imoto, 2018; Imoto & Shimauchi, 2016) in combination with automatic activity tracking. In such an approach, sound snippets of the acoustic scene are compared to a database and labeled accordingly. Paired with the localisation of a sound based on the stereo signal or possibly additional microphones, we may infer whether a sound was self-generated or not. However, we do not know whether the low sampling rates of acoustic features that we use is sufficient to capture all of these complexities; it will remain challenging when our goal is also to protect a persons’ privacy.

In conclusion, we showed again that we can record ear-EEG in everyday life and gain event-related potentials to discrete, well-defined acoustic events. In this study, we investigated the ERP response to experimentally presented sounds in everyday situations as well as naturally occurring sounds. We find clear ERPs for the experimentally presented sounds, but the ERPs to naturally occurring sounds are inconclusive. Additional information about natural sounds and the activity of participants in future investigations is necessary. Overall, our study provides evidence for the potential of beyond the lab studies using ear-EEG, but some challenges still need to be addressed.

## 5 Open Practices Statement

The EEG datasets are not publicly available as we consider them personalized data, but they are available from the corresponding author on reasonable request. The code used for the analyses and the experiment including the stimuli are available at https://doi.org/10.5281/zenodo.7361441.

## 6 Acknowledgement

This work was funded by the Deutsche Forschungsgemeinschaft (DFG, German Research Foundation) under the Emmy-Noether program - BL 1591/1-1 - Project ID 411333557. We thank our colleagues from the Neuropsychology Lab for the constructive discussions about this work and we thank Silvia Korte, Marc Rosenkranz, Manuela Jäger, Thorge Haupt and Arnd Meiser for their valuable comments on the manuscript. We thank Reiner Emkes and Sven Kissner for their technical support. We thank Katerina Limpitsouni for her illustrations found at undraw.co.

## 7 Competing Interests Statement

The authors have no conflict of interests to declare.

## Supplementary Material

**Figure 1:**
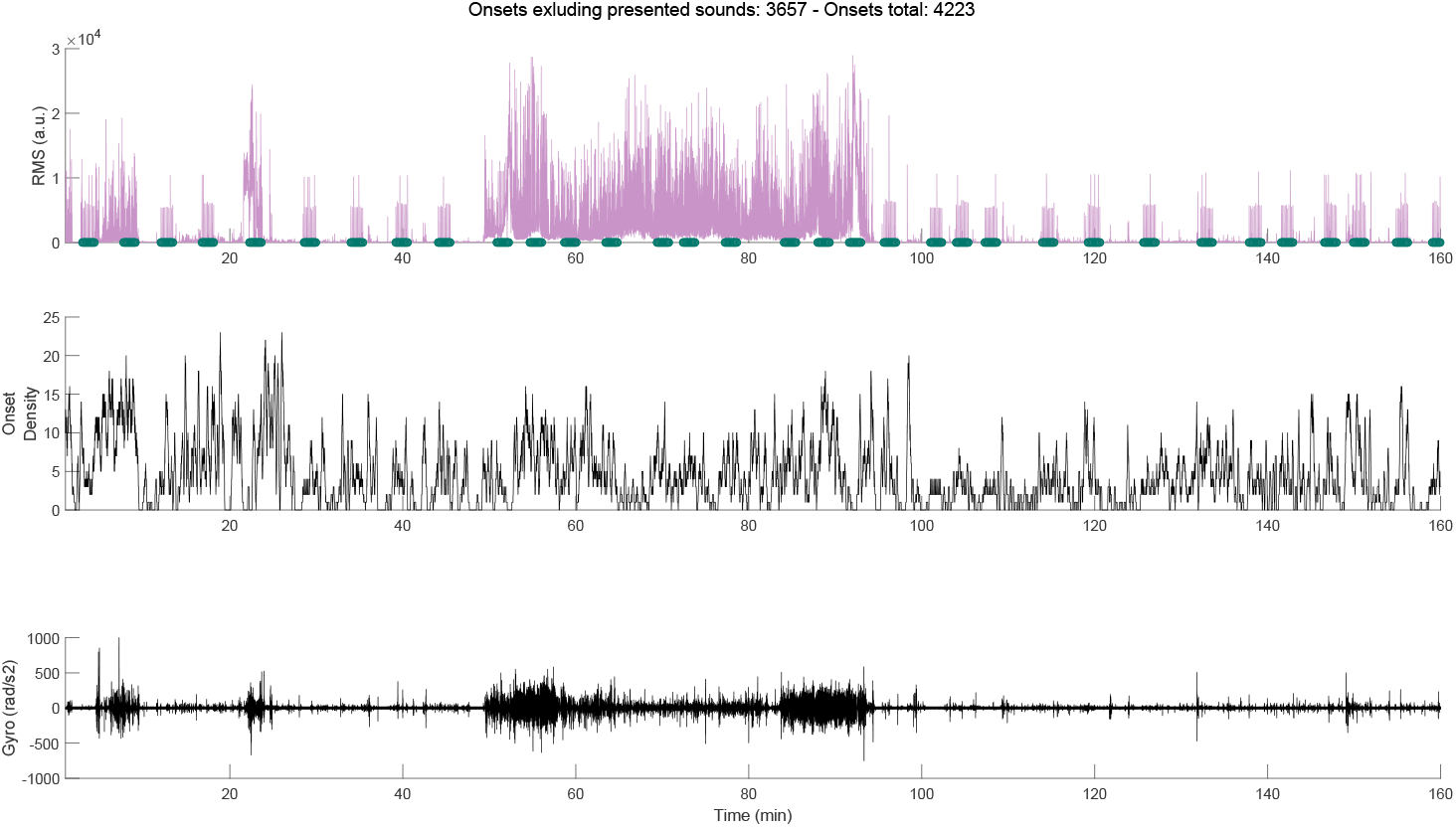
Time course of RMS, onset density, and gyro for participant 1. Green markers denote presented tones (sequences).

**Figure 2:**
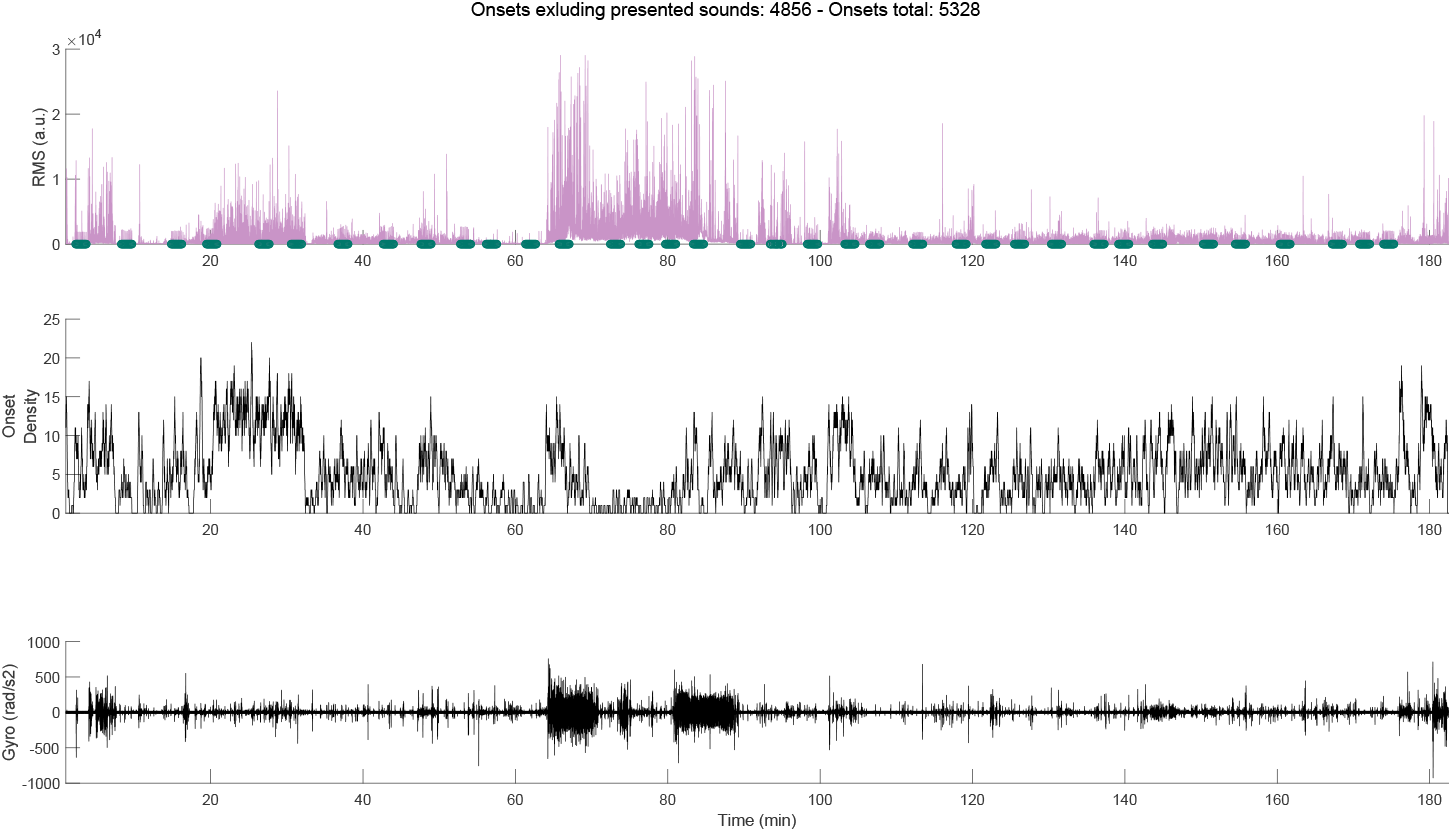
Time course of RMS, onset density, and gyro for participant 2. Green markers denote presented tones (sequences).

**Figure 3:**
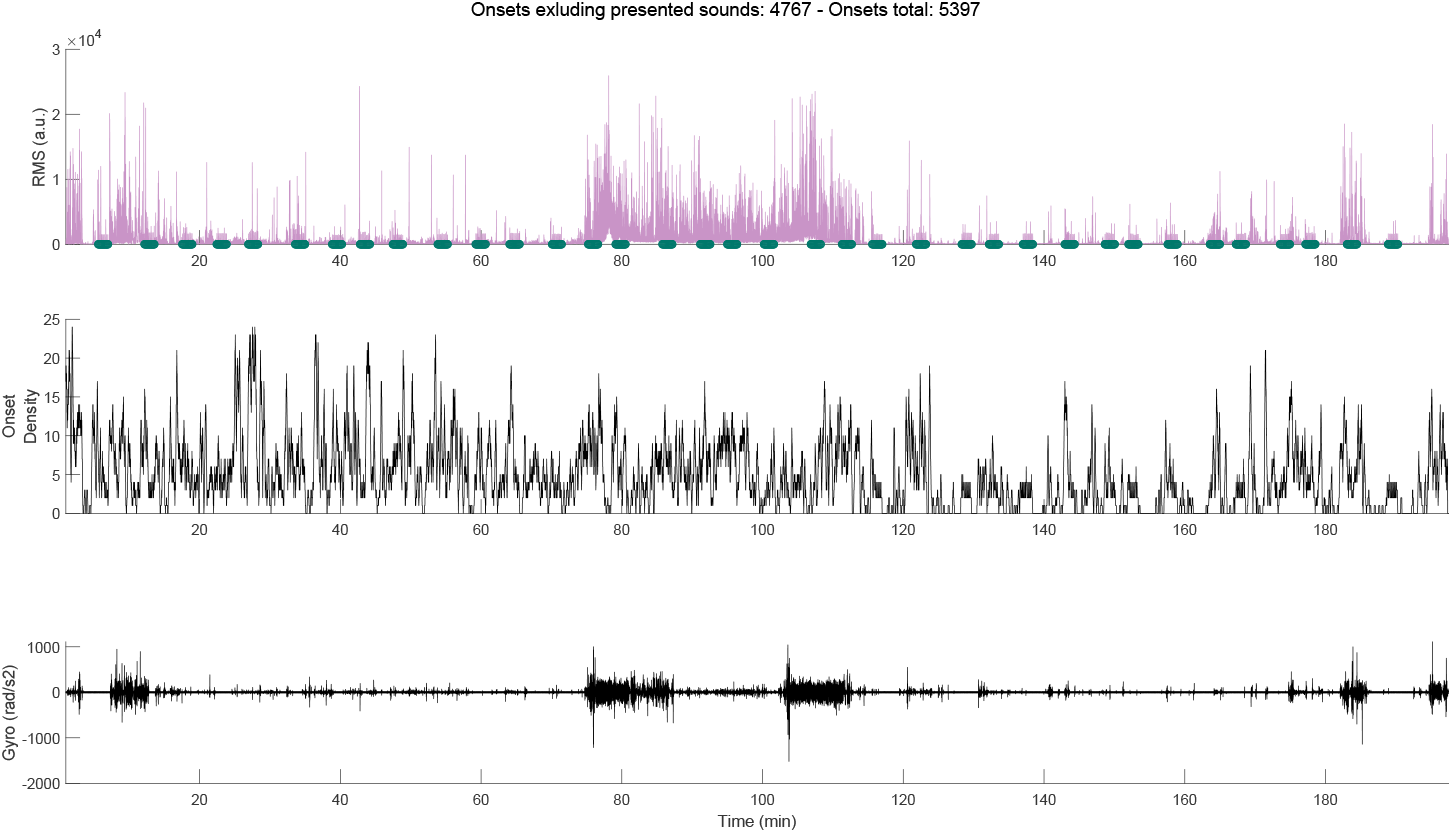
Time course of RMS, onset density, and gyro for participant 3. Green markers denote presented tones (sequences).

**Figure 4:**
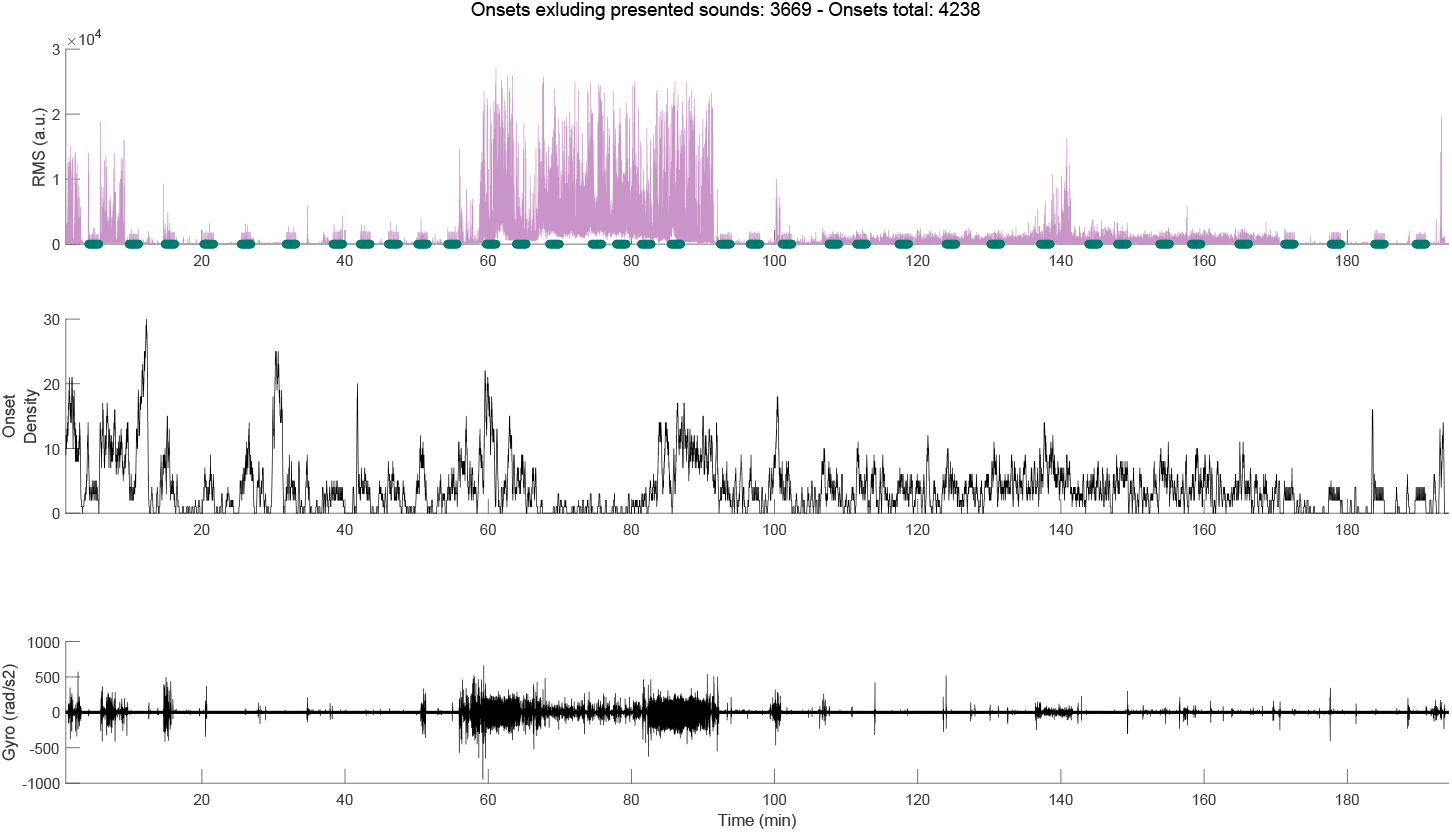
Time course of RMS, onset density, and gyro for participant 4. Green markers denote presented tones (sequences).

**Figure 5:**
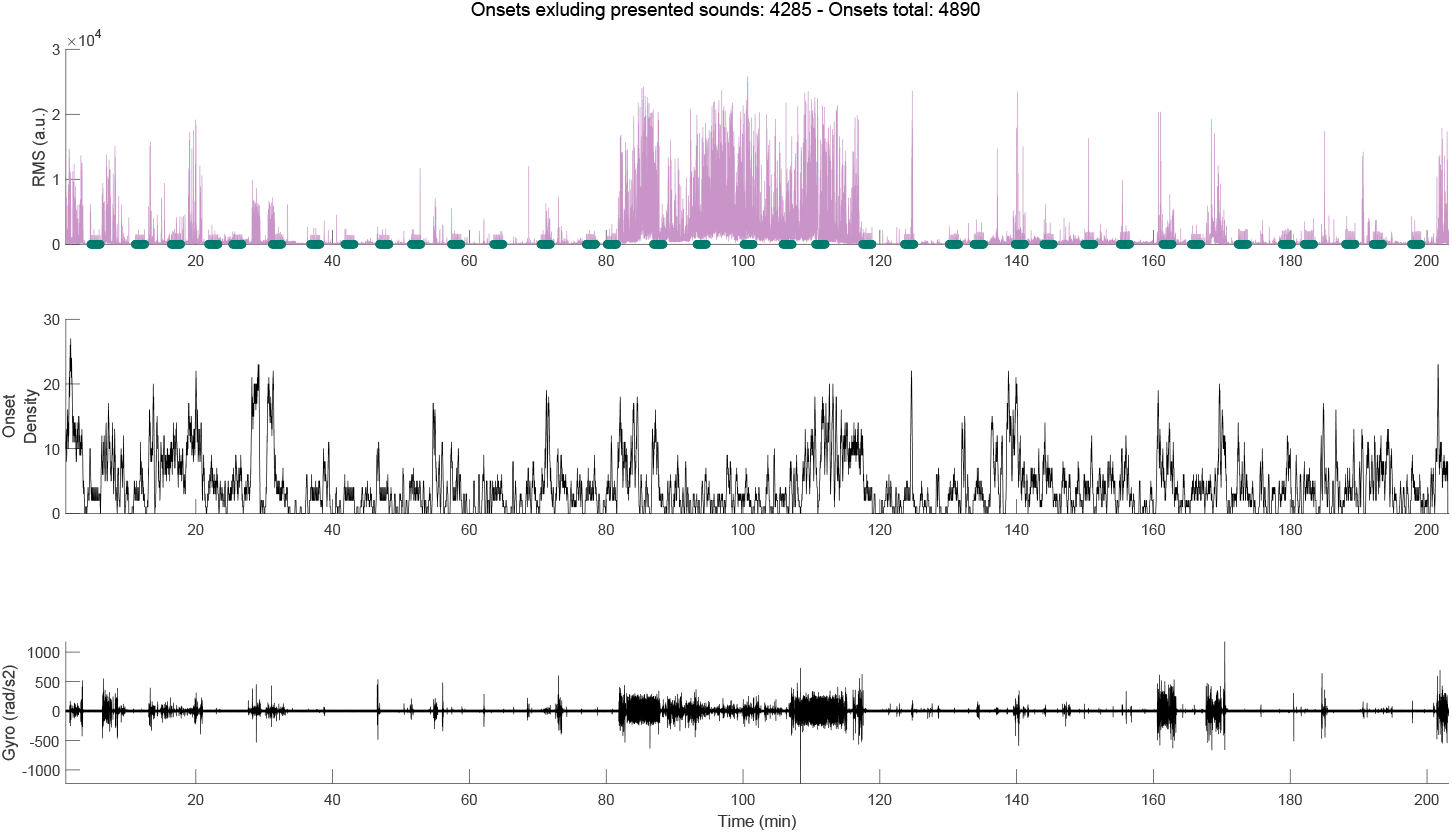
Time course of RMS, onset density, and gyro for participant 5. Green markers denote presented tones (sequences).

**Figure 6:**
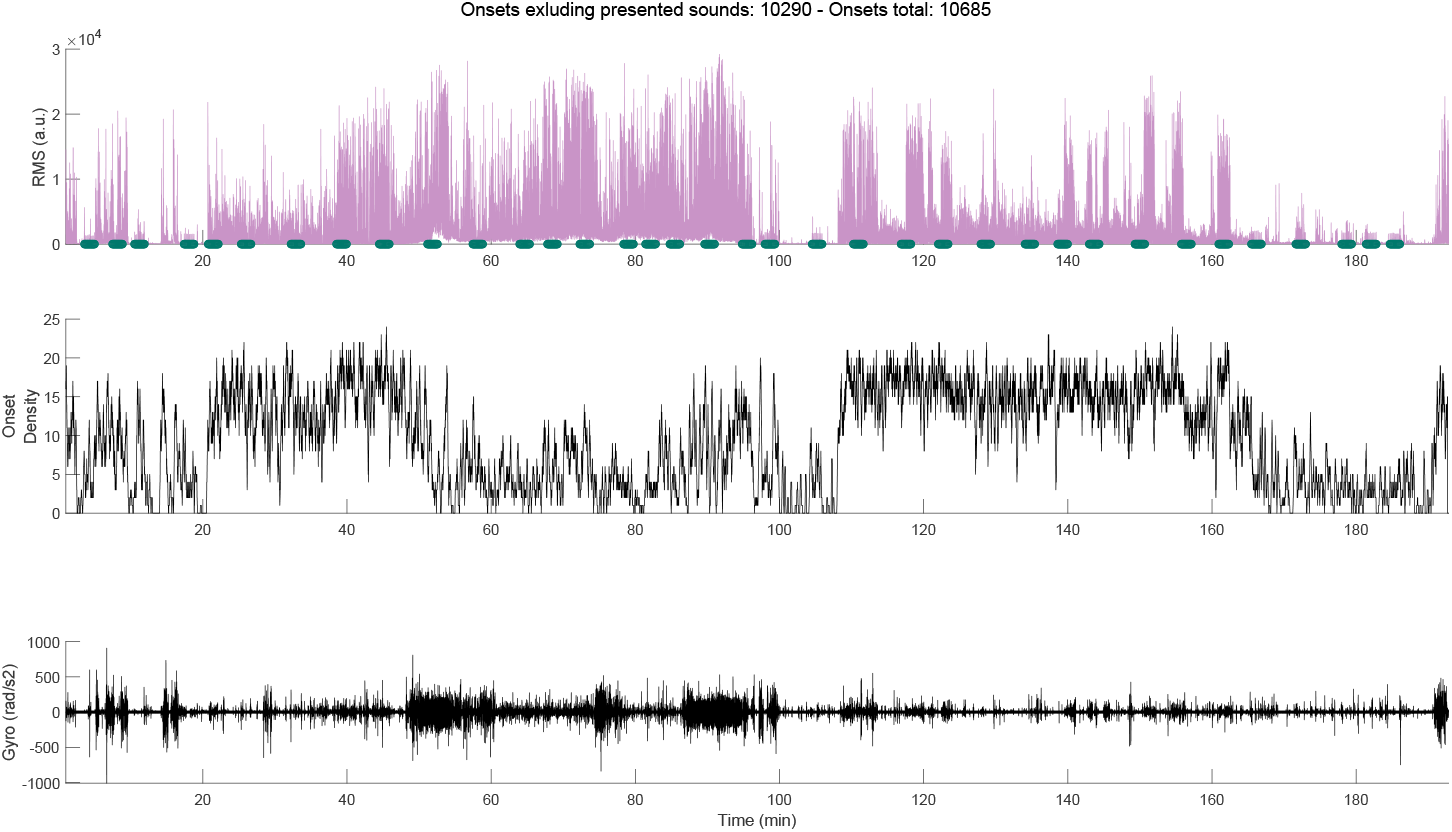
Time course of RMS, onset density, and gyro for participant 6. Green markers denote presented tones (sequences).

**Figure 7:**
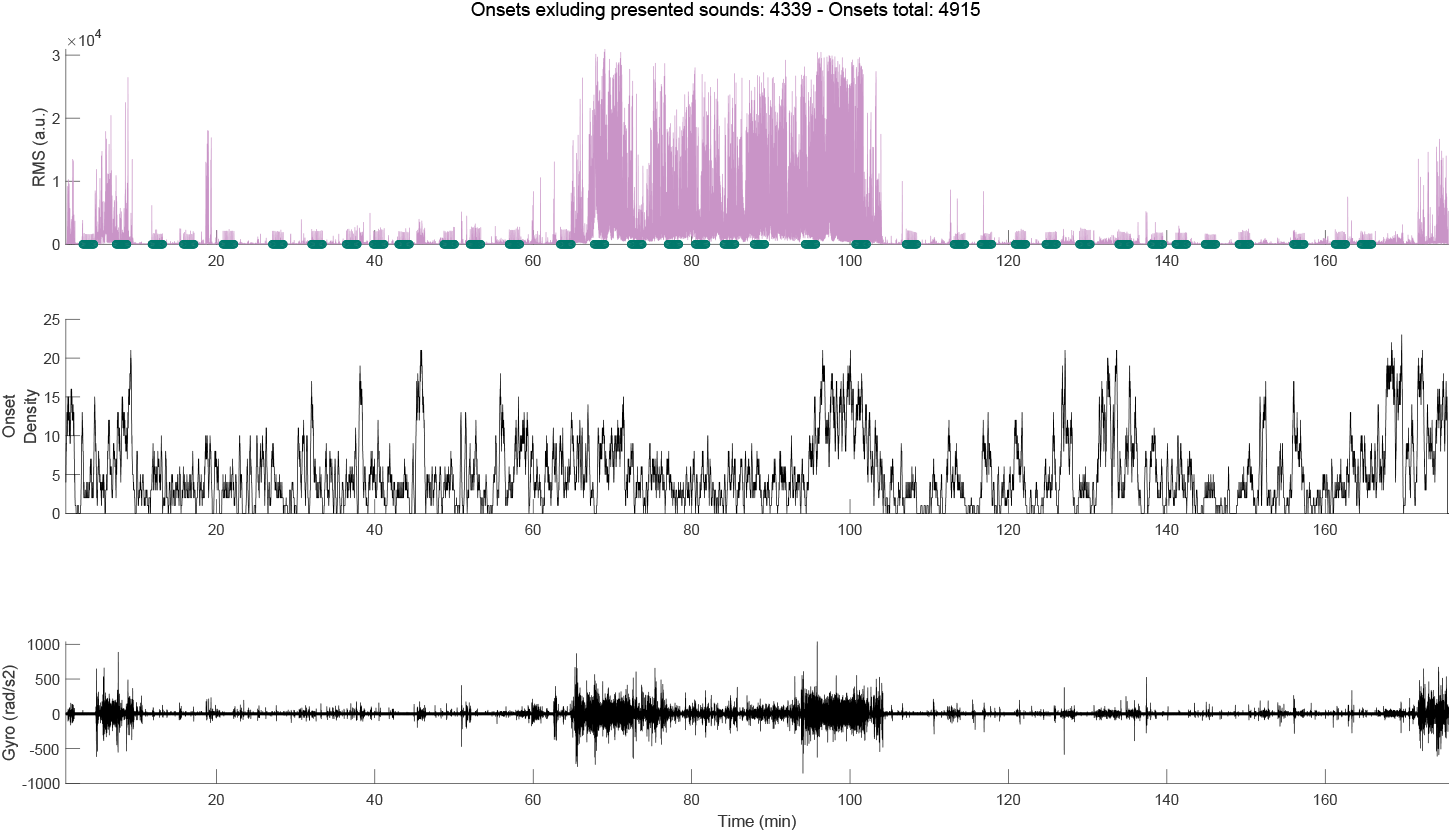
Time course of RMS, onset density, and gyro for participant 7. Green markers denote presented tones (sequences).

**Figure 8:**
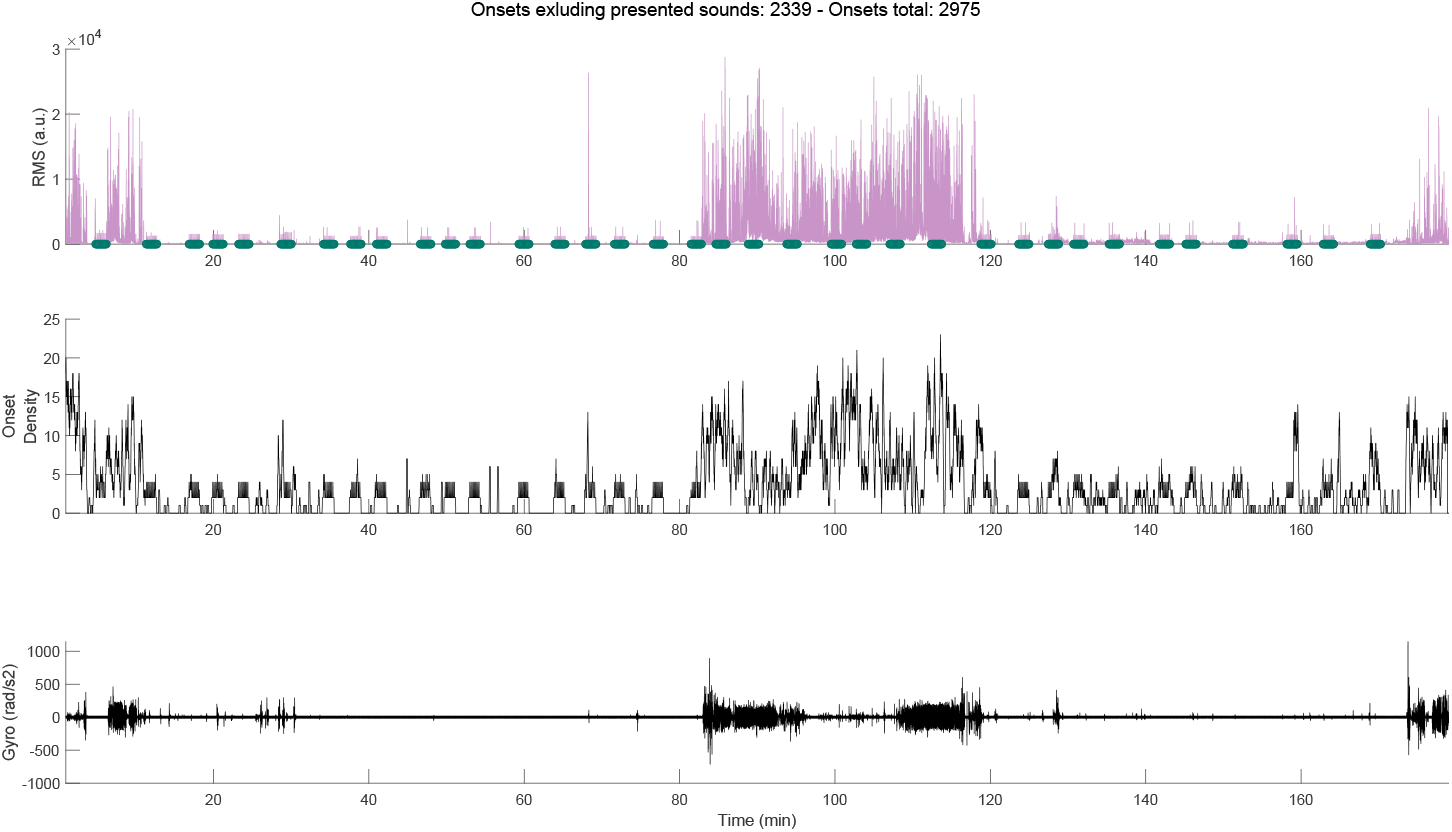
Time course of RMS, onset density, and gyro for participant 8. Green markers denote presented tones (sequences).

**Figure 9:**
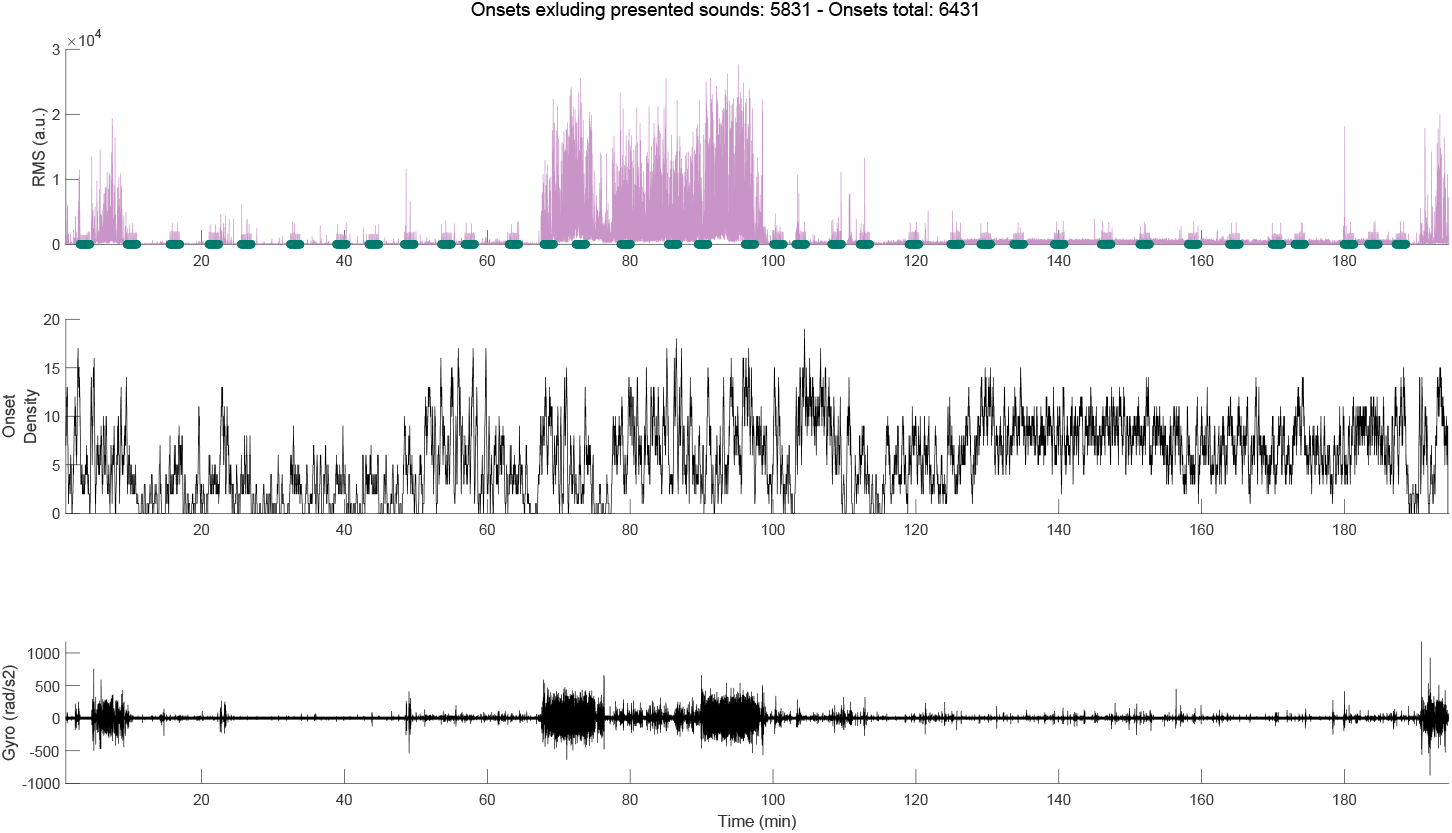
Time course of RMS, onset density, and gyro for participant 9. Green markers denote presented tones (sequences).

**Figure 10:**
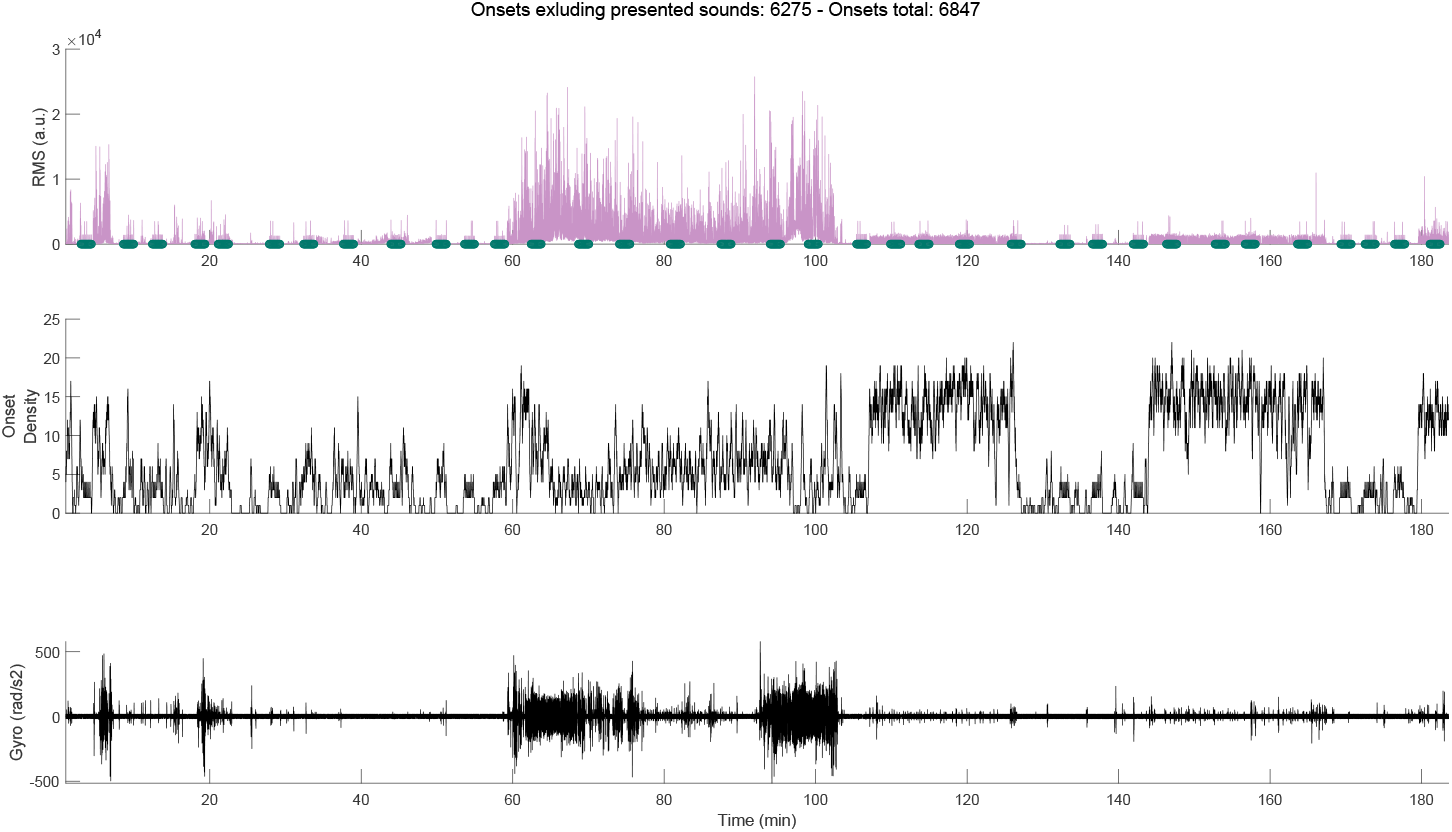
Time course of RMS, onset density, and gyro for participant 10. Green markers denote presented tones (sequences).

**Figure 11:**
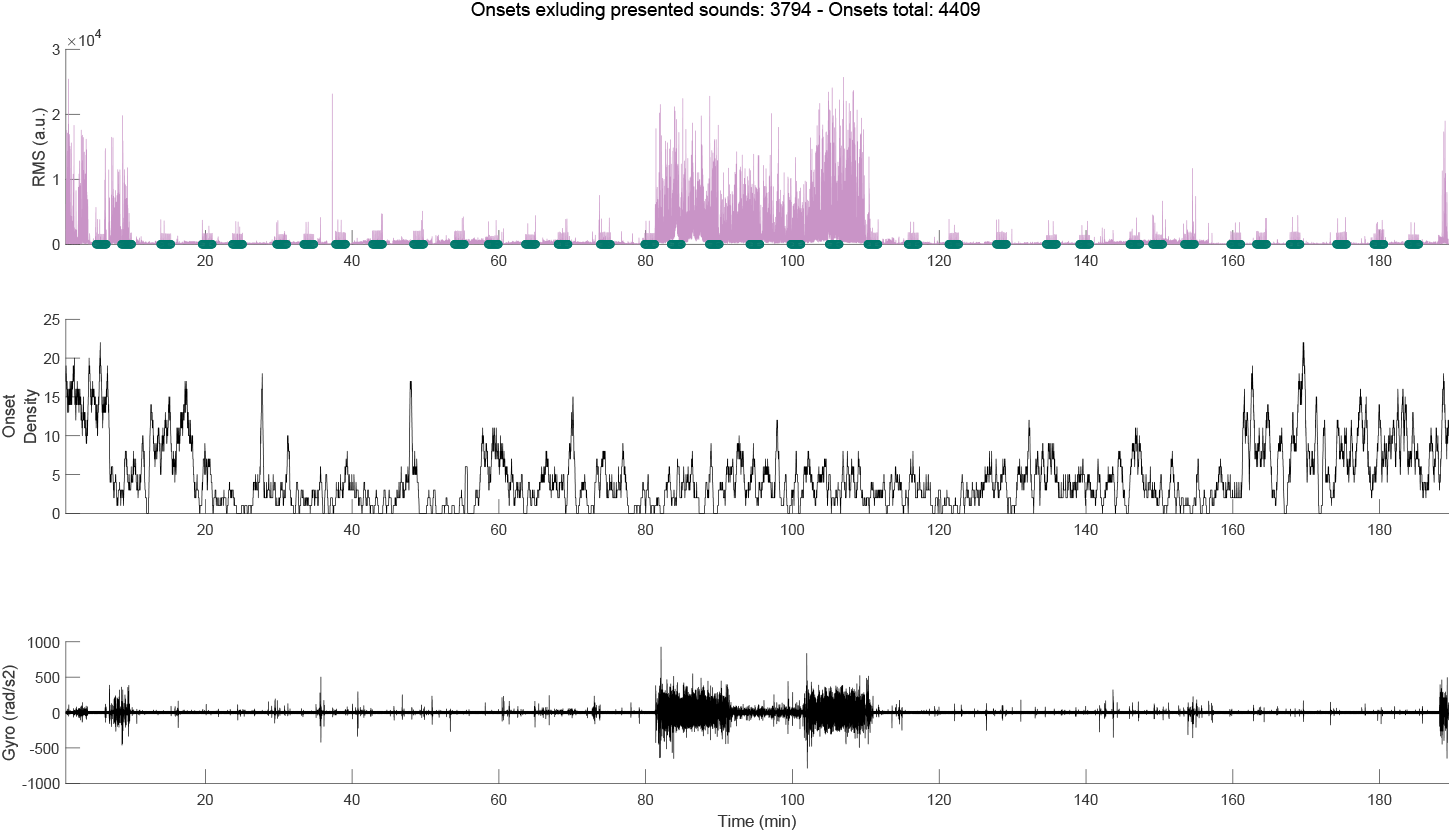
Time course of RMS, onset density, and gyro for participant 11. Green markers denote presented tones (sequences).

**Figure 12:**
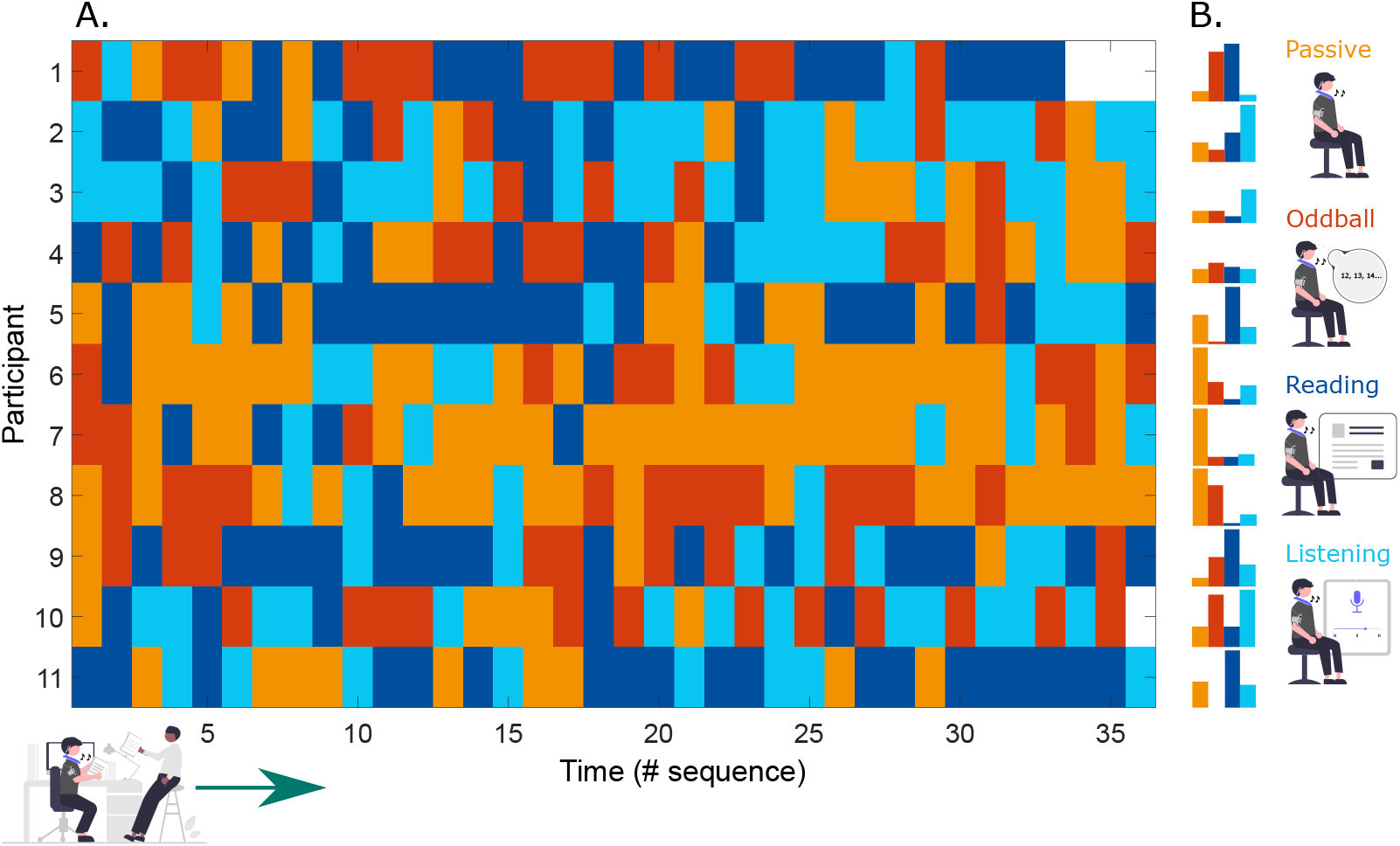
**A**. Results of the classification (n=11) of each sequence based on the lab reference conditions. This classifier was trained on all four lab conditions. Note that two participants have missing sequences at the end due to battery failure. **B**. The bars on the right visualize the percentage of each mode per participant. Ten-fold cross-validation was used to evaluate the classifier performance. The prediction error ranged from approximately 12% to 24% (M = 17.47%, SD= 4.15%)

**Table 1:** Linear mixed model for N1

| Fixed Effects | Model Summary |  |  |  |
| --- | --- | --- | --- | --- |
| | $\beta$ | SE | t-value | p-value |
| Intercept | -12.95 | 3.15 | -4.11 | <.001 |
| Background Noise | 7.17 | 3.58 | 2.01 | 0.046 |

| Random Effects |  |  |
| --- | --- | --- |
|  | Name | Standard Deviation |
| Participant | Intercept | 6.51 |

*Note:* The intercept is the predicted mean ERP amplitude ( μV) for low background noise. The beta for *Background Noise* is the predicted mean change in ERP amplitude when moving from low background noise to high background noise (i.e., the difference in amplitude between conditions).

**Table 2:** Linear mixed model for P1

| Fixed Effects | Model Summary |  |  |  |
| --- | --- | --- | --- | --- |
| | $\beta$ | SE | t-value | p-value |
| Intercept | 6.86 | 2.25 | 3.06 | <.01 |
| Background Noise | -3.80 | 3.04 | -1.25 | 0.21 |

| Random Effects |  |  |
| --- | --- | --- |
|  | Name | Standard Deviation |
| Participant | Intercept | 2.30 |

*Note:* The intercept is the predicted mean ERP amplitude ( μV) for low background noise. The beta for *Background Noise* is the predicted mean change in ERP amplitude when moving from low background noise to high background noise (i.e., the difference in amplitude between conditions).

**Table 3:** Linear model for P2

| Fixed Effects | Model Summary |  |  |  |
| --- | --- | --- | --- | --- |
| | $\beta$ | SE | t-value | p-value |
| Intercept | 13.34 | 2.30 | 5.82 | <.001 |
| Background Noise | 0.18 | 3.24 | 0.06 | 0.96 |

*Note:* The intercept is the predicted mean ERP amplitude ( $\mu V$ ) for low background noise. The beta for *Background Noise* is the predicted mean change in ERP amplitude when moving from low background noise to high background noise (i.e., the difference in amplitude between conditions).

**Table 4:** Linear model for P3

| Fixed Effects | Model Summary |  |  |  |
| --- | --- | --- | --- | --- |
| | $\beta$ | SE | t-value | p-value |
| Intercept | 11.57 | 2.32 | 4.98 | <.001 |
| Background Noise | -6.82 | 3.29 | -2.08 | 0.039 |

*Note:* The intercept is the predicted mean ERP amplitude ( $\mu V$ ) for low background noise. The beta for *Background Noise* is the predicted mean change in ERP amplitude when moving from low background noise to high background noise (i.e., the difference in amplitude between conditions).

## Notes

https://doi.org/10.5281/zenodo.7361441

### Competing Interest Statement

The authors have declared no competing interest.

